# Reintroduced native *Populus nigra* in restored floodplain reduces spread of exotic poplar species

**DOI:** 10.1101/2020.07.02.183699

**Authors:** An Vanden Broeck, Karen Cox, Alexander Van Braeckel, Sabrina Neyrinck, Nico De Regge, Kris Van Looy

## Abstract

Exotic *Populus* taxa pose a threat to the success of riparian forest restoration in floodplain areas. We evaluated the impact of exotic *Populus* taxa on softwood riparian forest development along the river Common Meuse after introducing native *Populus nigra* and after the re-establishment of the natural river dynamics. We sampled 154 poplar seedlings that spontaneously colonised restored habitat and assessed their taxonomy based on diagnostic chloroplast and nuclear microsatellite markers. Furthermore, by using a paternity analysis on 72 seedlings resulting from six open pollinated *P. nigra* females, we investigated natural hybridization between frequently planted cultivated poplars and native *P. nigra*. The majority of the poplar seedlings from the gravel banks analyzed where identified as *P. nigra*; only 2% of the sampled seedlings exhibited genes of exotic poplar species. Similarly, the majority of the seedlings from the open pollinated progenies were identified as *P. nigra*. For three seedlings (4%), paternity was assigned to a cultivar of *P*. x *canadensis*. Almost two decades after reintroducing *P. nigra*, the constitution of the seed and pollen pools changed in the study area in favour of reproduction of the native species and at the expense of the exotic poplar species. This study indicates that, although significant gene flow form exotic poplars is observed in European floodplains, restoration programmes of the native *P. nigra* can vigorously outcompete the exotic gene flows and strongly reduce the impact of exotic *Populus* taxa on the softwood riparian forest development.

## 2. Introduction

The restoration and protection of riparian forests is one of the key priorities in biodiversity conservation and climate change adaptation strategies (e.g. EU Biodiversity Strategy 2030, EU Floods Directive 2007/60/EC). Riparian forests are biodiversity hotspots and provide a range of ecosystem services including flood protection, prevention of bank erosion, thermal regulation by forest canopy cover and water quality protection (e.g. Van Looy et al. 2013). They are therefore necessary for the ecological functioning of riparian corridors and are recognized as an important part of the world’s natural capital.

After decades of ecological degradation caused by human activities, including impoundment and straightening of rivers, cutting off meanders, drainage of floodplains and deforestation for agriculture and plantations, several European countries have undertaken actions to restore natural river dynamics (e.g. Mansourian et al. 2019; Schindler et al. 2016), and the European Commission proposed a Biodiversity Strategy planning to restore at least 25,000 km of free-flowing rivers (UE COM Biodiversity Strategy 20/05/2020). Generally, restoration activities involve dyke relocation, lowering of river banks and floodplains to restore the floodplain habitat heterogeneity, an essential feature for the regeneration of riparian forests. However, many restoration projects are poorly documented and written records on the outcome of restoration projects are generally lacking (Palmer et al. 2007).

Revegetation of river margins is a widely applied technique for riparian forest restoration (Palmer et al. 2007). However, exotic, invasive alien species can significantly undermine efforts to protect and restore natural floodplain forests (Palmer et al. 2007). Exotic *Populus* species have been introduced into Europe as well as into the US and Canada for wood production in riparian landscapes and pose a threat to native *Populus* species. Poplar cultivars are frequently planted in monoclonal plantations in the vicinity of wild relatives and may contribute to a large extent to pollen and seed pools, thereby competing with the native relatives in colonizing restored habitat. Furthermore, intercrossability among *Populus* species is well-known (Eckenwalder 1984, 1996). In Europe, the native European black poplar (*Populus nigra* L.) (hereafter referred to as black poplar) is threatened by Euramerican (*P*. × *canadensis* Moench.), interamerican (*P*. × *generosa* Henry) hybrid poplar and black poplar varieties such as the male Lombardy poplar (*Populus nigra* cv. Italica Du Roi) (e.g. Arens et al. 1998; Cagelli and Lefèvre 1995; Chenault et al. 2011; Debeljak et al. 2015; Fossati et al. 2003; Ziegenhagen et al. 2008). Gene flow between cultivated and native poplar species may result in the reduction of the effective population size, genetic swamping and ultimately the replacement of populations of the native species (Arnold et al. 2001).

As a pioneer species, black poplar plays a key-role in the development of softwood forests in Europe. In the lower parts of the floodplain, black poplar colonises bare moist soil and river banks and is highly adapted to water dynamics and sediment movement (Debeljak et al. 2015; Van Looy 2006). Due to their rapid growth and strong flow resistance, black poplar together with species of *Salicaceae* (*Salix alba* and *S. viminalis*), contributes to raising bar and island levels by retaining sand and gravel (Van Looy 2006). Consequently, these softwood forest species pave the way for the development of the next forest succession stage, the hardwood forests. Unfortunately, the European black poplar is one of the most threatened tree species in Europe, mainly because of the loss of its natural alluvial habitats especially sand and gravel banks that allow for successful reproduction, and because of competition and introgression with exotic poplar species (Fossati et al. 2003; Meyer et al. 2018; Vanden Broeck et al. 2005; Ziegenhagen et al. 2008).

Here, we report on the impact of exotic *Populus* taxa at the initial stages of softwood riparian forest development of the river Meuse on the Dutch-Belgian border almost two decades after introducing black poplar and after the re-establishment of the natural river dynamics. To our knowledge, this is the first study reporting on the effect of re-introducing a native *Populus* species on gene flow of exotic relative species. The specific objectives of this study are (i) to study the taxonomy of the poplar seedlings that spontaneously colonised the river banks of the Common Meuse and (ii) to determine the frequency of natural hybridization events between male cultivated poplars (including *P. nigra* cv. Italica) and the female native black poplars in the study area. We hereto use a combination of diagnostic chloroplast and nuclear molecular markers that have been proven useful in identifying the taxonomy of *Populus* seedlings (Heinze 1998; Imbert and Lefèvre 2003; Ziegenhagen et al. 2008). We compare the results with a similar study performed in 1999 -2001 at the same site and before the restoration actions took place (Vanden Broeck et al. 2004). Finally, we discuss the implications for the restoration and conservation of softwood riparian forests with black poplar along European rivers.

## 3. Material & Methods

### 3.1 Study site

The area of interest was the riverside of the Common Meuse, a free-flowing middle course of the river Meuse, which forms for 45 km the border between Belgium and the Netherlands (Figure 1). Shipping and flow regulation are absent on this river stretch. The Common Meuse is a rain-fed river with a gravel-bed, a strong longitudinal gradient (0.45 m/km) and a wide alluvial plain (Van Looy et al. 2005). The Common Meuse valley consists of a gravel underground with a loamy alluvial cover. Traditionally, meadows were created in the floodplains. Cultivated poplars are frequently planted at the study site (Vanden Broeck et al. 2004). Large parts of the alluvial plain have been excavated for gravel mining, leaving large gravel pits or lowered floodplain zones (Van Looy et al. 2005). The extreme high water levels and floodings of 1993 and 1995, were the start of the development of a large scale, transboundary river restoration project for the Common Meuse; called ‘Living River Strategy’. The concept of this plan was to restore hydrodynamics and morphodynamics and related ecological characteristics in the primary river channel (Van Looy et al. 2006). The restoration activities were performed during the period 2005-2008 and included removal of dykes, excavation of the floodplain, channel widening and bank lowering. The restoration of the morphological activity (erosion / sedimentation rates) of the river reach resulted in a strong revitalisation of the elevation of bars and islands, creating suitable habitat for seedling recruitment and the restoration of softwood riparian forest with willow, poplar and ash communities (Van Braeckel 2007; Van Looy et al. 2008).

**Figure 1.**
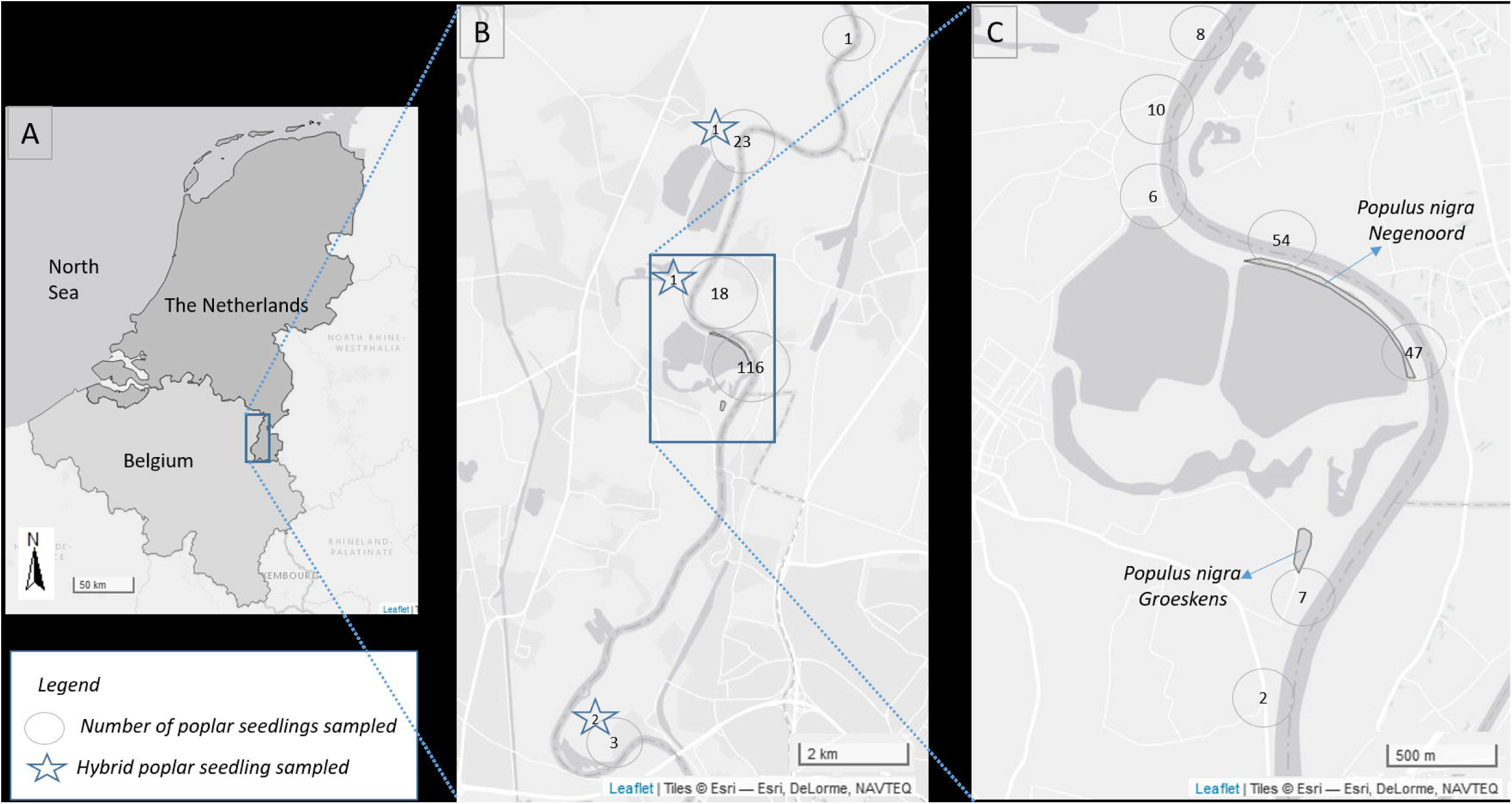
**A**. Location of the river Common Meuse. **B**. Map of the study site and sampling locations. Nearby sampled poplar seedlings are clustered and the total number of clustered samples is given in the circles. Stars indicate the position of hybrid poplar seedlings sampled. **C**. Location of the introduced *Populus nigra*.

### 3.2 Reinforcement of the Black poplar population

A reintroduction project of *P. nigra* was performed in the period 2002-2005. The aim was to create seed sources and thereby to promote the colonisation of native *P. nigra* in newly restored habitat. This was the follow-up of a study performed in 1999-2000 on the restoration potential and constraints of the development on softwood riparian forests along the Common Meuse (Vanden Broeck and Jochems 2002; Vanden Broeck et al. 2004). This former study emphasized the need for re-planting native *P. nigra* since natural populations were almost gone extinct in the study area and > 150 km further upstream in Northeast France (Vanden Broeck 2004). A few isolated relict trees of *P. nigra* in the study area suggested the presence of natural populations in the past. In contrast to the native poplar species, plantations of exotic poplars of *P*. x *canadensis* and *P. nigra* cv. Italica were common within the study area (Vanden Broeck et al. 2004).

*P. nigra* was successfully planted on two locations; location Groeskens (Dilsen-Stokkem, 51°01’01N, 05°46’02E) and location Negenoord (Dilsen-Stokkem, 51°01’36 N, 05°46’16E) on 0.77 ha and 2 ha, respectively. In total, 275 two-year old bare-root *P. nigra* trees were planted, with plant distances of 9 m x 9 m and 4 m x 8 m, in Groeskens and Negenoord, respectively. The distance from the centre of the plantings to the river bank was about 140 m for Groeskens and 5 m for Negenoord. The selection of the plant locations was based on the close connection to the river and on locations indicated close to potential sites for riparian forest development by a habitat model developed for the Common Meuse by Van Looy et al. (2005). The plant material originated form the national *P. nigra* gene bank collections at the Research Institute for Nature and Forest (INBO, Belgium; 57 genets) and at Alterra (The Netherlands; 21 genets) supplemented with 23 genets from a natural population located along the river Rhine in Germany (location Kühkopf, FA Groß-Gerau, 48° 484 56,62 N, 8° 7’ 38,60 O) and about 90 seedlings of controlled crosses performed at INBO with genotypes of *P. nigra* from the gene bank collection. All the plant material, was identified as pure *P. nigra* by molecular markers (Smulders et al. 2008b; Storme et al. 2001). Trees started to produce pollen and seeds, ten to fifteen years after planting.

### 3.3 Plant material sampled

#### 3.3.1 Poplar seedlings from the gravel banks

Poplar seedlings that naturally colonized the gravel banks of the natural floodplains of the river Common Meuse were sampled on the banks along a 28 km-river section. We collected leaf samples of 52 and 102 poplar seedlings in late summer of 2018 and 2019, respectively. The geographical location of each sampled seedling was recorded. In 2018, poplar seedlings were sampled randomly within the 28 km-section. In 2019, a more systematic survey was performed within part of the same route, of about 9 km along the riverside; each gravel bank was inspected by walking along the river. In case rejuvenation was present on a site, one to twenty-tree seedlings were sampled across the rejuvenation site (mean: 1.6), depending on the area of the colonized location. Only individuals that most likely originated from seed germination and not from vegetative reproduction by root suckers were sampled. One young leave was collected from each seedling and dried in silica gel for DNA-analyses. The estimated height of the majority of the sampled individuals was between 10 and 50 cm, corresponding with the seedling or establishing stage (2-3 years) of riparian forest development (Van Looy et al. 2005). The location along the river of the seedlings sampled is given in Figure 1. The poplar seedlings were visually identified by their leaf morphology. One young leave was collected from each seedling and dried in silica gel for DNA-analyses.

#### 3.3.2 Open pollinated progenies of *P. nigra*

*Populus* species are predominantly dioecious and thus obligatory outcrossers. Seeds of adult open pollinated (OP) European black poplar females located in the study area were collected in two successive years. In June 2018, seeds were collected at the location Groeskens from the lower branches of the crown of six European black poplars. In June 2019, seeds were collected on in total nine European black poplar females; six trees from location Groeskens and three trees from location Negenoord. Seeds were collected from mature catkins, sown in trays in the greenhouse (50% white peat / 50% black peat) within one week after collection. For the seeds collected in 2019, germination percentage was determined. One young leave was collected from each seedling and dried in silica gel for DNA-analyses. The number of seeds collected per open-pollinated *P. nigra* female is given in Table 1.

**Table 1.** The number of seeds collected, seed germination percentage and the number of seedlings analyzed per open pollinated *Populus nigra* female.

| Collection year | Location | P. nigra tree ID | # seeds | seed germination (%) | # seedlings analysed |
| --- | --- | --- | --- | --- | --- |
| 2018 | Groeskens | 2.16 | NA | NA | 28 |
| 2018 | Groeskens | 5.20 | NA | NA | 1 |
| 2019 | Negenoord | 1 | 160 | 2.50 | 4 |
| 2019 | Negenoord | 2 | 300 | 3.00 | 9 |
| 2019 | Negenoord | 3 | 300 | 5.67 | 17 |
| 2019 | Groeskens | 5.25 | 300 | 3.00 | 8 |
| 2019 | Groeskens | 5.20 | 76 | 6.58 | 5 |
| Total |  |  |  |  | 72 |

#### 3.3.3 Reference material

In Europe, poplar species frequently used in breeding programmes in order to produce hybrids are the European *P. nigra*, the North American cottonwoods *P. deltoides* W. Bartram ex Marshall. and *P. trichocarpa* Torr. & A. Gray., and to a lower extent *P. maximowiczii* Henry native to northeast Asia. The unit of cultivation and breeding in poplars is a clone, and individual cultivars are normally represented by a single clone. We included 76 reference samples with known taxonomic identity of frequently used pure poplar species and hybrids as positive controls for verifying the species-specificity of the molecular markers. Furthermore, reference samples of *P. nigra* and of frequently planted commercial male poplar cultivars were included as potential fathers in the paternity-analyses. Reference material was obtained from the clone collection of INBO located in Geraardsbergen, Belgium. The following taxa were included: section Aigeiros with *P. deltoides* (4 samples), *P. nigra* (18 samples), *P*. x *canadensis (P. deltoides* x *P. nigra*) (37 samples), section Tacamahaca with *P. trichocarpa* (4 samples), *P. trichocarpa* x *P. maximowiczii* (3 samples) and the intersectional hybrid; *P*. x *generosa* (*P. trichocarpa* x *P. deltoides)* (10 samples). The list of reference material is given in Supplementary Table S1.

### 3.4 Identification of hybrids

#### 3.4.1 DNA extraction

Total genomic DNA was extracted from the sampled leaves with the QuickPick Plant DNA kit (Bio-Nobile) using the MagRo 96-M robotic workstation (Bio-Nobile). For a subset of the samples (10%), the integrity of the DNA was assessed on 1% agarose gels. DNA quantification was performed with the Quant-iT PicoGreen dsDNA Assay Kit (Thermo Fisher Scientific, Massachusetts, USA) using a Synergy HT plate reader (BioTek, Vermont, USA).

#### 3.4.2 Chloroplast DNA marker

The chloroplast DNA (cpDNA) locus *trnDT* shows interspecific variation between *P. nigra, P. deltoides* and *P. trichocarpa* (Heinze 1998; Meyer et al. 2018). Because of its maternal inheritance and the female contribution of *P. deltoides* to the poplar clones frequently planted in Europe, this marker offers a tool to identify the maternal contribution of *P. deltoides* / *P*. x *canadensis, P. nigra* and *P. trichocarpa* in the poplar seedlings collected from the gravel banks (e.g. Csencsics and Holderegger 2016; Heinze 1997; Ziegenhagen et al. 2008). We used the conserved PCR primer pair trnD/trnT (Demesure et al. 1995) in combination with the restriction enzyme *Hinf*I. Genomic DNA (∼20 – 50 ng) was used as template in the PCR reactions. The PCR reaction conditions and restriction reactions were as described by Heinze (1998). Reaction products and a 100 bp ladder were analysed by agarose gel electrophoresis.

#### 3.4.3 Microsatellite analysis

We selected nine microsatellite loci that were found useful in former studies for the analysis of natural hybridization events between cultivated poplars and their wild relative *P. nigra* (Liesebach et al. 2010; Smulders et al. 2001; van der Schoot et al. 2000), and that were found reliable for clonal fingerprinting in several species and hybrids of the sections Aigeiros and Tacamahaca (Dayanandan et al. 1998; Liesebach et al. 2010; Rahman et al. 2000). All markers are unlinked (Cervera et al. 2001; Gaudet et al. 2008). According to former studies, five loci produce species-diagnostic alleles for *P. deltoides* and/or *P. trichocarpa:* PMGC14 *(Fossati et al. 2003; Liesebach et al. 2010)*, PMGC456 (Liesebach et al. 2010), PMGC2163 (Liesebach et al. 2010), WPMS09 (Fossati et al. 2003; Liesebach et al. 2010) and WPMS20 (Liesebach et al. 2010). SSR analysis was performed on the OP progenies, the seedlings from the gravel banks and the reference samples as described by Smulders et al. (2001) and van der Schoot et al. (2000). Ten samples were replicated twice starting from a second DNA-extract to calculate the genotyping error rate. PCR products were run on an ABI3500 Genetic Analyzer (Applied Biosystems) and genotypes were analysed with the GeneMapper v.6.0 software package. Samples with fewer than 8 scored loci were discarded from the data analysis. Details on the microsatellite loci are given in Table 2.

**Table 2.** Microsatellite loci used for taxonomic classification and paternity analysis.

|  | Locus | Linkage group | Repeat motif | Fragmentsize (bp) | Annealing Temperature (°C) | Reference |
| --- | --- | --- | --- | --- | --- | --- |
| 1 | PMGC_14 | XIII | CTT | 179 - 227 | 52 | PMGC |
| 2 | WPMS_05 | XII | GT | 263 - 291 | 52 | van der Schoot et al. (2000) |
| 3 | PMGC_2163 | X | GA | 198 - 220 | 52 | PMGC |
| 4 | PMGC_456 | II | GA | 80 - 136 | 52 | PMGC |
| 5 | WPMS_16 | VII | GTC | 128-167 | 52 | Smulders et al. (2001) |
| 6 | WPMS_09 | VI | GT | 244 -294 | 57 | Smulders et al. (2001) |
| 7 | PTR2 | IX | TGG | 207 -228 | 57 | Dayanandan et al. (1998) |
| 8 | WPMS_20 | XIII | TTCTGG | 224 - 242 | 57 | Smulders et al. (2001) |
| 9 | PTR7 | XII | (CT)5AT(CT) | 230 -250 | 57 | Rahman et al. (2002) |

#### 3.3.4 Data analysis

The maternal origin of the poplar seedlings sampled on the gravel banks was investigated using the cpDNA locus *trnDT*. The poplar seedlings from the gravel banks and the OP progenies were analysed for the presence of diagnostic alleles of *P. deltoides* and *P. trichocarpa* on the five diagnostic SSR loci. A diagnostic allele for *P. deltoides* in *P. nigra* maternal offspring indicates a backcrossed individual with *P*. x *canadensis* paternity as a result of a natural hybridization event between a cultivar of *P*. x *canadensis* and a native *P. nigra*.

We also performed a paternity analysis on the OP progenies using the microsatellite data to determine pollination events of frequently planted male poplar cultivars included in the reference samples. We used a likelihood-based approach implemented in the program CERVUS 3.0.7 (Marshall et al. 1998) with the reference samples described above (48; females excluded) as potential fathers, given a known mother. The paternity assignment technique used by CERVUS requires the knowledge of the total number of candidate males in the population and the proportion of the candidate males sampled for the simulation-based approach to assess the confidence of the assignments. For the simulation, we used the LOD distribution and we assumed 50 candidate fathers and a proportion of 0.3 candidate fathers sampled, allowing an error rate of 0.01 and allowing a mismatch between parent and offspring on one locus taking into account somatic mutations and null alleles.

## 4. Results

### 4.1 Diagnostic microsatellite alleles

The percentage missing data for the total microsatellite data was 0.07%. Replicated samples resulted in identical multilocus genotypes. For the seedlings sampled from the gravel banks, two multilocus genotypes were shared by two and three sampled seedlings, respectively. It is possible that some individuals were sampled twice, in 2018 and 2019. Only one genotype among these replicates was kept in the final dataset. The microsatellite data confirm the presence of species-diagnostic alleles reported in former studies by Liesebach et al. (2010) and Fossati et al. (2003) for PMGC14, PMGC456, PMGC2163 and WPMS09 but not for WPMS20. For PMGC14, the alleles 192 and 198 were typical for all reference clones of *P. deltoides* and for *P*. x *canadensis*, and the allele 200 was typical for all reference clones of *P. trichocarpa*. For WPMS09, allele 232 was present in the reference clones of *P. deltoides* and of *P*. x *canadensis*, except for 16 *P*. x *canadensis* reference clones that were homozygote for this locus. This can be explained by their genetic origin (i.e. F2 or back-cross clones) or by the presence of a null allele. For WPMS20, the characteristic allele for *P. deltoides* (allele 200) also frequently occurred in *P. nigra*, which was also reported by Liesebach et al. (2010). Furthermore, species-specific alleles were present at locus PMGC2163 (*P. deltoides*: allele 185, *P. trichocarpa*: alleles 199, 200) and locus PMGC456 (*P. deltoides*: allele 82, *P. trichocarpa*: alleles 93, 109, *P. nigra*: allele 75).

### 4.2 Poplar seedlings from the gravel banks

From the 154 seedlings sampled on the gravel banks, 135 showed a clear banding pattern for the cpDNA locus *trnDT*. From the latter, 133 (98.5%) poplar seedlings showed a *P. nigra*-specific *trnDT* marker-fragment (∼880 bp). These seedlings also showed a *P. nigra*-like leaf morphology. The typical gene variant of *P. nigra* observed on the maternal inherited diagnostic cpDNA locus *trnDT* indicated that they originated from a female *P. nigra*. Furthermore, two (1.5%) seedlings showed a *P. trichocarpa*-specific cpDNA fragment (∼1000bp). These two samples with a *P. trichocarpa*-like leave morphology were collected on an upstream distance of 6 km from the nearest *P. nigra*. No seedlings were detected with a fragment length specific of *P. deltoides* (∼1135bp). The results of the diagnostic microsatellite alleles confirmed the presence of species-specific *P. trichocarpa* alleles at loci PMGC147, PMGC2163 and PMGC456 for the two seedlings with a *P. trichocarpa*-specific *trnDT* marker-fragment. For one of these two seedlings, species-specific alleles of *P. trichocarpa* were homozygous, suggesting that this seedling could have originated from two *P. trichocarpa* or *P*. x *generosa* cultivars. Furthermore, two more seedlings sampled in the field showed species-specific alleles of *P. deltoides* in heterozygous state at three and four diagnostic loci, respectively. Given their *P. nigra*-specific *trnDT* marker-fragment, they likely originated from a *P. nigra* x *P*. x *canadensis* back-cross. Summarised, four (2%) out of the 154 young poplars analysed from the gravel banks exhibited genes of exotic, cultivated poplar species.

### 4.3 Identification of hybrids in open pollinated progenies

Viable seedlings were obtained from six and five black poplar trees in 2018 and 2019, respectively. The mother trees presented only two different genets. The mean germination percentage for the 1492 seeds sown in 2019 was low (2.9%; range of 0% to 6.6%). In total, we obtained 74 viable seedlings of which 72 were successfully genotyped (Table 1). We detected no alleles species-specific to *P. deltoides* or *P. trichocarpa* in the 72 analysed seedlings from the OP *P. nigra* progenies, suggesting the absence of backcrossed individuals with *P*. x *canadensis* or *P*. x *generosa* as potential fathers.

However, the paternity analysis revealed *P*. x *canadensis* a highly likely father for three seedlings from the OP progeny from seed harvested in 2018 on one mother tree (ID 2.16) at location Groeskens. Paternity was determined with 95% confidence for 27 (38%) seedlings. When using a relaxed confidence level of 80%, paternity was assigned to 48 (67%) seedlings. For 24 (33%) seedlings, the paternity remained unresolved. For one seedling paternity was assigned to *P*. x *canadensis* cv. Serotina de Champagne using a relaxed confidence level and with no paternity-offspring mismatches. Also for a second seedling from the same mother tree, *P*. x *canadensis* cv. Serotina de Champagne was the most likely father with a mismatch for only one locus (PMGC14), possibly due to a somatic mutation. A third seeding from the same OP progeny showed *P*. x *canadensis* cv. Serotina as the most likely father, although mismatch was observed at two loci (PMGC14, PTR07). The other determined paternities were assigned to trees of *P. nigra*. No offspring was assigned to the male cultivar *P. nigra* cv. Italica, nor to the nearby planted male *P*. x *canadensis* cv. Robusta or to the reference cultivars of *P*. x *generosa*.

## 5. Discussion

The results of this study illustrate the importance of in situ reinforcement of black poplar to reduce the potential impact of exotic poplar cultivars in European floodplains in case natural river dynamic processes are restored and adult black poplar trees are scarce. The numbers of young poplar seedlings that colonized the gravel banks of the river Meuse exceeded 100 fold the initial number of seedlings observed by Vanden Broeck et al. (2004). Additionally the number of seeds and seedlings with exotic poplar genes were diminished to a very low proportion compared to the situation before the restoration.

### 5.1 Limited interspecific matings

The majority (150, 98%) of the poplar seedlings from the gravel banks analyzed where identified as ‘pure’ *P. nigra* seedlings; the typical gene variant of *P. nigra* observed in the maternal inherited diagnostic cpDNA locus *trnDT* indicated that they originated from a female *P. nigra*. They could not originate from cultivated first-generation (F1) *P*. x *canadensis* hybrids, as cultivated clones of *P*. x *canadensis* are the result from artificial crosses between a female *P. deltoides* and a male *P. nigra*, thus carrying the *P. deltoides* haplotype (Heinze 1998; Zsuffa 1974). Furthermore, the absence of microsatellite alleles of *P. deltoides* or *P. trichocarpa* suggests that they were fathered by *P. nigra*. Only a small number (four; 2%) of young poplars analyzed from the gravel banks exhibited genes of exotic, cultivated poplars. Of the latter, two seedlings showed *P. trichocarpa* genes while the other two resulted from a first-generation back-cross (BC1) of a male *P*. x *canadensis* and a female *P. nigra*. No gene variants of exotic poplar species were detected in the OP progenies. This was largely confirmed by the paternity analyses, except for three seedlings (4%) of which two were likely fathered by *P*. x *canadensis* cv. Serotina de Champagne and one seedling by *P*. x *canadensis* cv. Serotina. It must be noted that these two cultivars are clearly different and not related, although their names suggest otherwise. They show different alleles at seven out of the nine microsatellite loci genotyped and also show differences in leaf morphology as was noted by Broekhuizen (1960). No paternity was assigned to the male cultivar *P. nigra* cv. Italica (Lombardy poplar), one of the most planted poplar cultivars. In Belgium, the Lombardy poplar generally flowers before the majority of the native black poplars (Vanden Broeck et al. 2003a), which may not be the case in more Southern regions (Chenault et al. 2011). Absence of flowering synchrony may explain why no paternity was assigned to this cultivar or to the nearby planted male *P*. x *canadensis* cv. Robusta in this study.

To our knowledge, this is the first study combining diagnostic chloroplast and nuclear molecular markers with a paternity analysis to detect interspecific hybridization events between native black poplar and poplar cultivars. The probability to detect gene variants of *P. deltoides* using four diagnostic nuclear microsatellite loci and according to Mendelian rules in first-backcross generations (BC1) is 93.75%, but decreases to 68.36% for second back-cross generations (BC2) (see Bialozyt et al. 2012). This could explain why gene variants of *P. deltoides* remained undetected in three seedlings from the OP progenies to which a cultivar of *P*. x *canadensis* was assigned as the most likely father. In this study, the paternity analysis increased the reliability to identify further generation back-cross events in the OP progenies. However, the power of a paternity analysis depends on the potential male parents included in the analysis and the assumptions made on the proportion of potential fathers sampled. It is possible that some second back-cross events remained undetected because of these sampling limitations.

### 5.2 Pollen and seed pools affecting exotic gene flow

The contrast in the number of interspecific mating events before and after the river and vegetation restoration activities is remarkable. In 1999 - 2001, the majority of the seedlings sampled on the gravel banks (27/29; 93%) showed a *P. trichocarpa* fragment at the cpDNA locus *trnDT* and/or species-specific alleles of *P. deltoides* (Vanden Broeck et al. 2004). In contrast to this study, gene variants of exotic poplar species were also detected in the majority of the seedlings (32/34; 94%) of the OP progenies of an isolated female *P. nigra* in the study area in 1999 - 2001 (Vanden Broeck et al. 2004). At that time, *P. nigra* was extremely rare in the study area as a result of habitat reduction and fragmentation; the native species was only represented by a few old trees, in contrast with the widely planted poplar cultivars (Vanden Broeck et al. 2004). The strong decrease in population size combined with the habitat fragmentation of the native *P. nigra* increased the opportunities for contact with cultivated poplar clones and provided at that time excellent opportunities for the production of hybrid seeds and seedlings. Poplars are prolific seed producers; one tree can produce over 50 million of seeds in a single season (Oecd 2000). Nevertheless, poplar hybrids are generally characterized by reduced fertility relative to parental species, with significant lower pollen production and seed viability in F1-hybrids (Stettler et al. 1996). It is therefore likely that the seed production and seed viability of *P. nigra* outperformed those of the cultivated poplars. Also, pollen load composition and size could have been an important process in limiting the production of hybrid progeny. The low frequencies of interspecific mating relative to intraspecific mating using a pollen mixture of different taxa in controlled crosses suggest a greater competitive ability for conspecific pollen (Benetka et al. 2002; Gaget et al. 1989; Rajora 1989; Vanden Broeck et al. 2003b). As a result, the relative fertilization success of a pollen tree depends upon the taxon constitution of the pollen mix and thus on the number and taxon of the producing pollen trees in the study area.

In this study, the few seedlings showing exotic poplar genes were located at distances of > 1 km from the nearest reintroduction sites of black poplar. This indicates that hybrid poplars still reproduce in the study area, in particular on sites located further away from the reforestation sites of black poplar. Establishing more seed sources of black poplar over the river stretch could further reduce hybrid reproductive success along the Common Meuse. Although conspecific siring of female *P*. x *canadensis* is enhanced by the presence of pollen of black poplar (Vanden Broeck et al. 2012), the higher local density of male and female black poplars appears to guarantee the establishment of mainly pure black poplar seedlings nearby the parental stands. Considering the possibility of long distance gene flow (Imbert and Lefèvre 2003), investigations of interspecific mating events would ideally cover the whole river stretch where natural habitat is restored.

### 5.3 Conservation implications

The results of this study illustrate the importance of in situ reinforcement of black poplar populations to reduce the potential impact of exotic poplar cultivars along European floodplains where natural river dynamic processes are restored and adult black poplar trees are scarce. The potential risks for gene introgression from exotic poplars into black poplar is indeed highly variable between geographic locations and river sites, largely depending on habitat fragmentation, flowering synchrony and the composition of the pollen and seed pool at the specific site. While interspecific gene flow seems to be absent in large natural populations of black poplar, like in France along the rivers Loire, Garonne and Drôme (Imbert and Lefèvre 2003), significant gene flow from exotic poplars in riparian floodplains is reported along the rivers Rhine and the Elbe in Germany (Meyer et al. 2018; Ziegenhagen et al. 2008), the Morava River in the Czech Republic {Pospiskova, 2006 #832}, the Ticino river in Italy (Fossati et al. 2003), the Rhine river in The Netherlands (Arens et al. 1998; Smulders et al. 2008a) and the rivers Thur and Reuss in Switzerland (Csencsics and Holderegger 2016). This case study along the Common Meuse, demonstrates that reintroduction and reinforcement of native black poplar populations can reverse the situation from high to low risk of introgression with cultivated poplars on a relatively short time scale. By changing the composition of the seed and pollen pool, black poplar nowadays has a competitive advantage compared to the exotic poplars in colonizing the restored river banks along the Common Meuse.

Hundreds of young black poplar seedlings colonized the gravel banks. With an estimated age for the majority of the seedlings of two to three years old, they survived the germination stage and currently present the establishment stage of softwood forest development with the development of dense thickets (Van Looy 2006). The survival of poplar seedlings depends on the combination of many factors broadly related to the availability of water and oxygen (Hughes et al. 2001). As a result of the critical conditions required by seedlings, successful establishment only occurs in some years, and within well-defined elevational bands (Hughes et al. 2001). Furthermore, established seedlings at lower river levels are frequently subject to later removal or damage by drought, flooding, sediment burial or mechanical disturbance (Hughes et al. 2001; Jacquemyn et al. 2006). Seedling survival should therefore be monitored over larger time scales to evaluate the success of this restoration project and to predict riparian forest development along the river Common Meuse. Management and conservation plans should include further monitoring of the seedling colonization and survival, the effective population size and the introgression of foreign genes in black poplar across multiple generations. Also, the vegetative spread of cultivated hybrids, which was not the focus of this study, should be taken into consideration in predicting the development of riparian forest ecosystems. The data can then be used for the construction of habitat models to identify occupied or potential habitat and to predict riparian forest development in the restored floodplain area (Debeljak et al. 2015; Van Looy et al. 2005).

## 6. Conclusions

Almost two decades after the reintroduction of black poplar along the river Common Meuse, the constitution of the seed and pollen pool changed in the study area thereby benefitting the reproduction of the native species at the expense of the reproduction capacity of the exotic poplar species. Increasing the density of male and female black poplars appears to guarantee the establishment of mainly pure black poplar seedlings nearby the parental stands. Nevertheless, exotic poplar species still reproduce at larger distances from the revegetation sites and colonize restored habitat thereby competing for resources with the native species. Therefore, increasing the number of native black poplar seed sources along the river stretch will help the native black poplar to maintain its competitive advantage over the exotic poplars and will contribute considerably to the success of riparian forest development along the Common Meuse.

## 7. Author statements

AVdB, KVL: conceptualization; AVdB: methodology, formal analysis, writing - original draft; KC, AVB, KVL: writing - review & editing; SN, NDR: methodology, resources. All authors have read and approved the final version.

## 9. Acknowledgements

The authors gratefully acknowledge the river managers Herman Gielen and Joke Verstraelen for their enthusiasm and support. The authors also thank Wim De Clercq and Jemp Peeters for the field assistance, and Marc Schouppe for the technical assistance in the greenhouse.

## 11. Data profile

Data will be made available after acceptance of the manuscript for publication by the data repository DRYAD.

## 15. Supplementary material

**<u>Table S1</u>**. List of *Populus* species and poplar cultivars with known taxonomic identity used as reference samples. References to the cultivars were obtained from the International Register of Poplar Cultivars (IPC – FAO)

